# Once-daily feeding is associated with better health in companion dogs: Results from the Dog Aging Project

**DOI:** 10.1101/2021.11.08.467616

**Authors:** Emily E. Bray, Zihan Zheng, M. Katherine Tolbert, Brianah M. McCoy, Dog Aging Project Consortium, Matt Kaeberlein, Kathleen F. Kerr

## Abstract

A variety of diets have been studied for possible anti-aging effects. In particular, studies of intermittent fasting and time-restricted feeding in laboratory rodents have found evidence of beneficial health outcomes. Companion dogs represent a unique opportunity to study diet in a large mammal that shares human environments. The Dog Aging Project has been collecting data on thousands of companion dogs of all different ages, sizes, and breeds since 2019. We leveraged this diverse cross-sectional dataset to investigate associations between feeding frequency and cognitive function (*n* = 10,474) as well as nine broad categories of health conditions (*n* = 24,238). Controlling for sex, age, breed, and other potential confounders, we found that dogs fed once daily rather than more frequently had lower mean scores on a cognitive dysfunction scale, and lower odds of having gastrointestinal, dental, orthopedic, kidney/urinary, and liver/pancreas disorders. Therefore, we find that once-daily feeding is associated with better health in multiple domains. Future research with longitudinal data can provide stronger evidence for a possible causal effect of feeding frequency on health in companion dogs.

## Introduction

For nearly a century, caloric restriction has been known to extend lifespan and delay age-associated pathology in laboratory animals [1–5]. More recently, in both animals and humans, a variety of alternative “anti-aging” diet modalities have been described which are providing new mechanistic insights and potential clinical applications [6]. These diets include intermittent fasting [7, 8], fasting mimicking diets [9, 10], ketogenic diets [11–15], protein or essential amino acid restriction [16, 17], and time-restricted feeding [18–20].

These diets have been most extensively studied in rodents in controlled laboratory settings, due to ease of administering diets on a specific schedule and an enhanced ability to tease apart the mechanisms through which they act. Time-restricted feeding studies in rodents suggest improvements in several metabolic parameters, including glucose and insulin homeostasis, energy expenditure, hepatic pathology, resistance to different obesogenic diets, and improved circadian rhythm maintenance during aging [21–23]. In one study, mice who experienced time-restricted feeding demonstrated an 11% extension in lifespan [18]. Additionally, several studies demonstrate that caloric restriction and intermittent fasting play a protective role in maintaining and enhancing cognitive function, including memory and spatial learning [24–28]. It remains unclear, however, whether these benefits in laboratory animals are generally due to reduced caloric intake or meal frequency or both [6].

Despite mainstream popularization in humans of several of these diets, the beneficial health effects of time-restricted feeding outside of a laboratory setting are less clear. In some human studies, only mild improvements in body composition and cardiovascular risk factors were detected [29], even when subjects also reduced their daily caloric intake [30]. In other studies, detrimental effects on glucose homeostasis were observed with time-restricted feeding [31]. Finally, while some studies have found potential cognitive benefits, especially for memory in older adults [32, 33], other studies have shown no effect of fasting on cognition [34, 35].

Companion dogs provide a potentially powerful animal model to elucidate the relationship between diet and age-related health outcomes [36]. Having co-evolved alongside people for thousands of years [37], companion dogs share human environments, experience similar diseases, and receive similar medical care. Once-daily feeding in dogs serves as a natural model for the intermittent fasting/time-restricted feeding protocols currently being studied both in preclinical rodent models and in human trials [38].

The Dog Aging Project is a large-scale research initiative following thousands of companion dogs over their lifetimes to better understand how biology, lifestyle, and environment impact healthy aging [39]. Participating owners report annually on a variety of aspects related to their dog’s diet, primary and secondary activities, social and physical environments, medications, and health conditions. In the current study, we used cross-sectional data collected in the first year of the Dog Aging Project to ask if feeding frequency is associated with cognitive function and health conditions. Specifically, we hypothesized that dogs fed once-a-day would display lower rates of physical health issues and better cognitive scores compared to dogs fed more frequently.

## Methods

### Subjects

All dogs had been recruited to join the Dog Aging Project (DAP) via mainstream media, social media, or word of mouth. Their owners then completed the relevant online surveys between December 26, 2019 and December 31, 2020 [40]. Study data were collected and managed using REDCap electronic data capture tools hosted through the DAP [41, 42]. REDCap (Research Electronic Data Capture) is a secure, web-based software platform designed to support data capture for research studies, providing 1) an intuitive interface for validated data capture; 2) audit trails for tracking data manipulation and export procedures; 3) automated export procedures for seamless data downloads to common statistical packages; and 4) procedures for data integration and interoperability with external sources.

### Instruments

The first survey that participants completed was the Health and Life Experience Survey (HLES), which collects information regarding dog demographics, physical activity, environment, dog behavior, diet, medications and preventives, health status, and owner demographics. In the current investigation we were principally interested in feeding frequency and health status, and we identified *a priori* health conditions that could plausibly be affected by feeding frequency. After completing HLES, all participants were offered the opportunity to complete the Canine Social and Learned Behavior Survey (CSLB), which measures cognitive function. The CSLB, renamed by the DAP, is the same as the validated Canine Cognitive Dysfunction Rating Scale (CCDR) [43], with only a handful of minor wording changes. The Canine Cognitive Dysfunction Rating Scale was presented to participants as the Canine Social and Learned Behavior Survey to avoid the negative connotations of the phrase ‘cognitive dysfunction. This instrument asks owners to indicate the frequency with which their dogs exhibit behaviors indicative of dementia (i.e., disengagement from social activity; difficulty in navigation, searching, and recognition). Based on owner responses, dogs receive a score that ranges from 16 to 80, where higher scores are indicative of worse cognitive function.

During the study time period, 27,541 DAP participants completed HLES, and 20,096 DAP participants completed CSLB.

### Ethical Note

The University of Washington IRB deemed that recruitment of dog owners for the Dog Aging Project, and the administration and content of the DAP Health and Life Experience Survey (HLES), are human subjects research that qualifies for Category 2 exempt status (IRB ID no. 5988, effective 10/30/2018). No interactions between researchers and privately owned dogs occurred; therefore, IACUC oversight was not required.

### Inclusion/Exclusion Criteria

Given that meal frequency is adjusted as puppies mature, we specified age inclusion as 1 ≤ age < 18 years for all health outcomes. For the CSLB outcome, we specified age inclusion as 6 ≤ age < 18 years, as 6 years is the youngest age indicated in the literature where signs of cognitive decline can start to appear in dogs [44–46]. Less than 5% of dogs in the DAP are intact and these dogs were excluded, as well as dogs (<1%) whose owners reported that their diet was “not at all consistent.” Thus, in our final sample, all dogs were spayed or neutered due to exclusion criteria, and slightly less than half of dogs were male. About one-fifth of dogs received daily or more frequent omega-3 or other fatty acid supplementation in their diet.

We studied health conditions that were reported in the nine broad categories on HLES that could plausibly be affected by feeding frequency: dental or oral disease, skin disorders, orthopedic disorders, gastrointestinal disorders, cancer or tumors, kidney or urinary disorders, cardiac disorders, neurological disorders, and liver or pancreas disorders. The other broad categories of health conditions reported in HLES were not analyzed because they were either based on temporary situational and/or environmental factors and thus unlikely to be associated with feeding frequency (e.g., trauma, ingesting toxic substances, infectious or parasitic disorders); were infrequently reported and thus had a very small sample size (less than 3.5% of the total sample; e.g., respiratory disorders, endocrine disorders, reproductive system disorders, immune-mediated disorders, and hematopoietic disorders); or there was no compelling rationale as to why feeding frequency would affect them (e.g., ear, nose, and throat disorders, eye disorders).

For the health categories examined in this investigation, all participants were assigned a binary score (affected/unaffected). Dogs were considered ‘affected’ if their owner reported them to have at least one relevant condition within a given category. However, we did not consider any congenital health outcomes as ‘affected’: since animals were born with these conditions, their feeding regimen was by definition instituted after onset and could therefore not have affected the development of the condition. Similarly, disorders linked to transient situational factors, including infectious diseases and trauma, were not considered as ‘affected,’ as the circumstantial nature of these instances made them unlikely to be affected by feeding frequency.

See Supplementary Information 1 for details of all inclusion/exclusion criteria, as well as the specific conditions that qualified a dog as affected within the dental or oral disease, skin disorders, orthopedic disorders, gastrointestinal disorders, cancer or tumors, kidney or urinary disorders, cardiac disorders, neurological disorders, and liver or pancreas disorders categories.

After applying exclusion criteria, the final sample consisted of responses from 24,238 HLES surveys and 10,474 CSLB surveys. The CSLB was always completed at least one week after completion of HLES. Most participants in the final sample (88%) completed CSLB within 3 months of completing HLES and always within a year (range: 7 to 352 days, average: 47 days).

### Explanatory variables

We analyzed feeding frequency as a binary exposure, comparing dogs fed once-daily to dogs fed more frequently. Specifically, owners were asked “How many times per day is your dog fed?” The dogs of owners who answered “Once” were sorted into the once-daily category, whereas the dogs of owners who answered “Twice”, “Three or more”, or “Free fed (filling up bowl when empty or always having food available)” were sorted into the fed-more-frequently category. In all analyses, 8% of the total sample were fed once daily (Table 1a and Table 1b).

**Table 1a.** Characteristics of 10,474 Dogs in CSLB Analysis (76 pure breeds included).

| Explanatory variable | n (%) or mean (SD) |
| --- | --- |
| **Age** |  |
| 6-9.9 years | 4956 (47%) |
| 10-18 (not including 18) years | 5518 (53%) |
| **Sex** |  |
| Male, castrated | 5042 (48%) |
| Female, spayed | 5432 (52%) |
| **Body Size & Weight** |  |
| Small (<10 kg) | 2673 (26%) |
| Middle (10-29.9 kg) | 5064 (48%) |
| Large (30-44.9 kg) | 2389 (23%) |
| Giant (≥45 kg) | 348 (3%) |
| **Breed** |  |
| Mixed-breed dog | 5885 (56%) |
| Labrador Retriever | 616 (6%) |
| Golden Retriever | 463 (4%) |
| German Shepherd | 192 (2%) |
| Dachshund | 159 (2%) |
| Australian Shepherd | 149 (1%) |
| Poodle (Standard) (≥13.6 kg) | 130 (1%) |
| Border Collie | 124 (1%) |
| Chihuahua | 107 (1%) |
| Beagle | 105 (1%) |
| Shih Tzu | 93 (1%) |
| Other purebred dogs<sup>1</sup> | 2451 (23%) |
| **Feeding Frequency** |  |
| Fed once per day | 860 (8%) |
| **Fatty Acid Supplement** |  |
| Given daily or more often | 2252 (22%) |
| **Cognitive Activity** |  |
| Primary or Secondary Activity<br>requires training | 1759 (17%) |
| **CSLB** |  |
| Score | 36.4 (5.3) |

<sup>1</sup> See Supplementary Information 4 for full list and numbers of purebred dogs included in the CSLB analysis.

**Table 1b.** Characteristics of 24,238 Dogs in Analysis of Health Outcomes (100 pure breeds included).

| Explanatory variable | n (%) |
| --- | --- |
| **Age** |  |
| 1-1.9 years | 1672 (7%) |
| 2-5.9 years | 7721 (32%) |
| 6-9.9 years | 7082 (29%) |
| 10-18 (not including 18) years | 7763 (32%) |
| **Sex** |  |
| Male, castrated | 11853 (49%) |
| Female, spayed | 12385 (51%) |
| **Body Size & Weight** |  |
| Small (<10 kg) | 5510 (23%) |
| Middle (10-29.9 kg) | 12267 (51%) |
| Large (30-44.9 kg) | 5504 (23%) |
| Giant (≥45 kg) | 957 (4%) |
| **Breed** |  |
| Mixed-breed dog | 13308 (55%) |
| Labrador retriever | 1467 (6%) |
| Golden retriever | 1199 (5%) |
| German Shepherd | 502 (2%) |
| Australian Shepherd | 373 (2%) |
| Poodle (Standard) (≥13.6 kg) | 306 (1%) |
| Dachshund | 305 (1%) |
| Border Collie | 257 (1%) |
| Chihuahua | 205 (1%) |
| Beagle | 187 (1%) |
| Pug | 180 (1%) |
| Other purebred dogs<sup>2</sup> | 5949 (25%) |
| **Feeding Frequency** |  |
| Fed once per day | 1884 (8%) |
| **Fatty Acid Supplement** |  |
| Given daily or more often | 4383 (18%) |
| **Health Outcomes** |  |
| Dental/Oral | 6414 (26%) |
| Skin | 5619 (23%) |
| Orthopedic | 4270 (18%) |
| Gastrointestinal | 2429 (10%) |
| Cancer | 1906 (8%) |
| Kidney/Urinary | 1740 (7%) |
| Cardiac | 1243 (5%) |
| Neurological | 972 (4%) |
| Liver/Pancreas | 673 (3%) |

<sup>2</sup> See Supplementary Information 5 for full list and numbers of purebred dogs included in the analysis of health outcomes.

In our analyses, we adjusted for sex (spayed female or neutered male), age, breed for purebred dogs and body size (as captured by weight) for mixed breed dogs. We also adjusted for whether the owner reported daily omega-3 (or other fatty acid) diet supplementation for all analyses except for dental/oral disorders and liver/pancreas, as there is evidence in the literature that fatty acids can have beneficial effects on cognitive function, skin, cardiac, gastrointestinal, renal, orthopedic, and neoplastic outcomes [47–55]. For analysis of CSLB, we additionally adjusted for two factors that are thought to affect cognitive function: physical activity level [56] and whether the dog has a history of training (according to the dog’s primary or secondary activity indicated by the owner; e.g., show dogs, service dogs, and dogs trained for field trials vs. pets/companion dogs; see Supplementary Information 2 for full details) [57].

We adjusted for the breed of purebred dogs as a categorical variable. After inspecting the distribution of weight by breed, we subdivided Standard poodles into two breeds for the analysis, large poodles (weight ≥ 13.6 kg (30 lb)) and small poodles (weight < 13.6 kg (30 lb)). Although there are over 200 breeds represented in DAP data, for each analysis we only included breeds that had at least one exposed and one unexposed dog because breeds without variance in the exposure cannot inform the exposure-outcome association. We also restricted our analyses to breeds with at least 10 dogs meeting inclusion criteria. These restrictions reduced the number of breeds to 76 breeds for the CSLB analysis and 100 breeds for the analyses of health outcomes.

### Statistical Methods

All statistical analyses were carried out in R v.4.0.3 [58]. Age was flexibly modeled using natural splines with interior knots at 7, 10, and 14 years for CSLB analysis and interior knots at 2, 7, and 13 years for health outcomes [59]. Weight was similarly modeled using natural splines with interior knots at 14, 48, and 79 lbs. In each instance, interior knots are at approximately the 10^th^, 50^th^, and 90^th^ percentile of each variable. We explored more elaborate adjustment models (e.g., 4 or 5 interior knots), but these were not supported by metrics such as AIC and examination of some results suggested overfitting.

To adjust for physical activity, we performed principal component analysis on three HLES-reported activity variables: lifestyle activity level (reported as not active, moderately active, or very active over the past year), average activity intensity level (reported as low: walking, medium: jogging, or vigorous: sprinting, such as playing fetch or frisbee), and average daily time spent physically active (reported in hours and minutes). Parallel analysis recommended retaining one principal component. This principal component captured 52% of the variance, and we used the loadings onto the first principal component as a physical activity score (PA-score). We adjusted for PA-score using natural splines with interior knots at approximately the 10^th^, 50^th^, and 90^th^ percentiles.

Mixed breed dogs were included as a separate category of breed, and we adjusted for body size, as measured by weight, for mixed breed dogs only by constructing a variable *weight*\**MB* where *weight* is the dog’s weight and *MB* = 1 for mixed breed dogs and *MB* = 0 for purebred dogs. This is analogous to grouping mixed breed dogs by weight and including each group as a breed, except that our approach uses continuous weight information.

We used linear regression for analysis of CSLB and logistic regression for analysis of all health outcomes. For linear regression, the large number of parameters in the model due to 76 breeds does not cause statistical issues. However, large models are problematic for logistic regression when using conventional maximum likelihood model-fitting [60]. Therefore, we fit the logistic models using a conditional likelihood, where the conditioning was on the breed categories. This approach allowed breed to be in the model without necessitating the estimation of 100 breed parameters. Due to the large size of the data set, maximizing the exact conditional likelihood was not computationally feasible, and we used the Efron approximation. We investigated the fidelity of the approximation with follow-up analyses (reported in Supplementary Information 3) on mixed breed dogs plus dogs from the 10 most common breeds and fitting the model with ordinary logistic regression. The 10 most common breeds were Australian shepherd, beagle, border collie, chihuahua, dachshund, German shepherd, golden retriever, Labrador retriever, poodles (large), and pugs (Table 1b). We also treated these analyses as secondary analyses to assess the robustness of our findings. We used robust standard error estimates for all regression analyses. All hypothesis tests were two-sided and we did not adjust for multiple comparisons.

## Results

In the CSLB analysis (*n* = 10,474 dogs), 56% of dogs were mixed breed with the remaining dogs belonging to 76 breeds (Table 1a and Supplementary Information 4). Dogs fed once per day had, on average, a 0.62 point lower CSLB score than dogs fed more than once per day (95%: 0.27, 0.97; *p* < 0.001), adjusting for age, sex, weight (for mixed breed dogs), breed (for purebred dogs), cognitive activity, physical activity level, and fatty acid supplementation (Figure 1; see Supplementary Information 3 for the full model). This effect size of 0.62 points is roughly the same difference in mean CSLB score between 11- and 7-year-old dogs.

**Figure 1:**
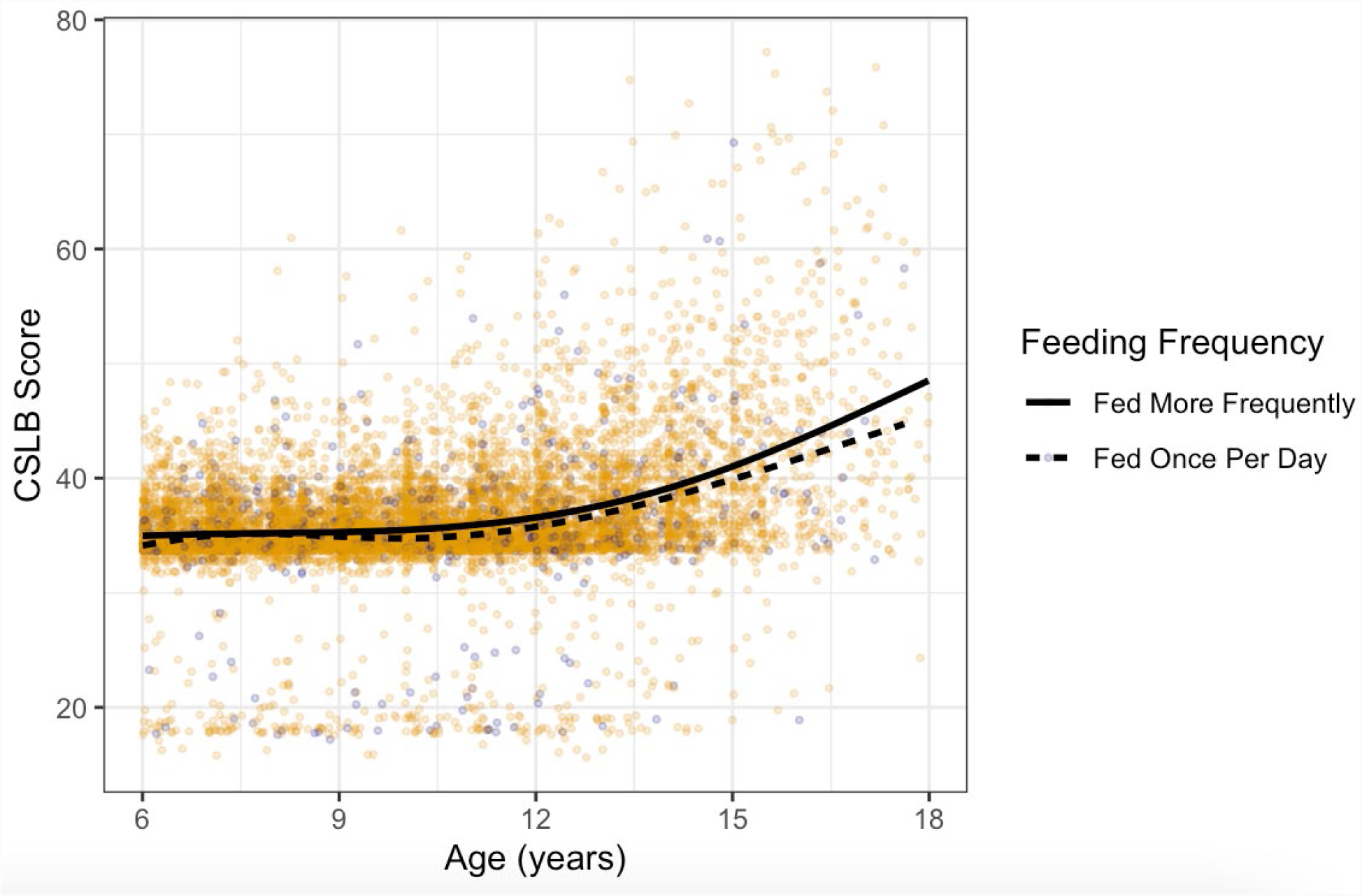
Scatterplot of CSLB Scores vs. Age with Superimposed Trend Lines. Darker points represent dogs fed once daily and other points represent dogs fed more frequently. Trend lines were constructed separately for the two groups using natural splines. Dogs fed once daily have slightly lower mean CSLB score at all ages (6 ≤ age < 18 years).

In analyses of health conditions (*n* = 24,238 dogs), 55% of dogs were of mixed breed and the remaining dogs belonged to 100 breeds (Table 1b and Supplementary Information 5). For five of nine health conditions analyzed, we found evidence that being fed once per day vs. more often is associated with lower odds of having the health condition (Figure 2; Table 2). Adjusted odds ratios were less than one and statistically significant for gastrointestinal, dental/oral, orthopedic, kidney/urinary, and liver/pancreas health conditions. Adjusted odds ratios were also less than one for the remaining four categories of health conditions (cardiac, skin, neurological, cancer), but not statistically significant (see Supplementary Information 3 for detailed model results). Results were similar in secondary analyses including only mixed breed dogs and dogs from the ten most common breeds (see Supplementary Information 3 for full report).

**Figure 2:**
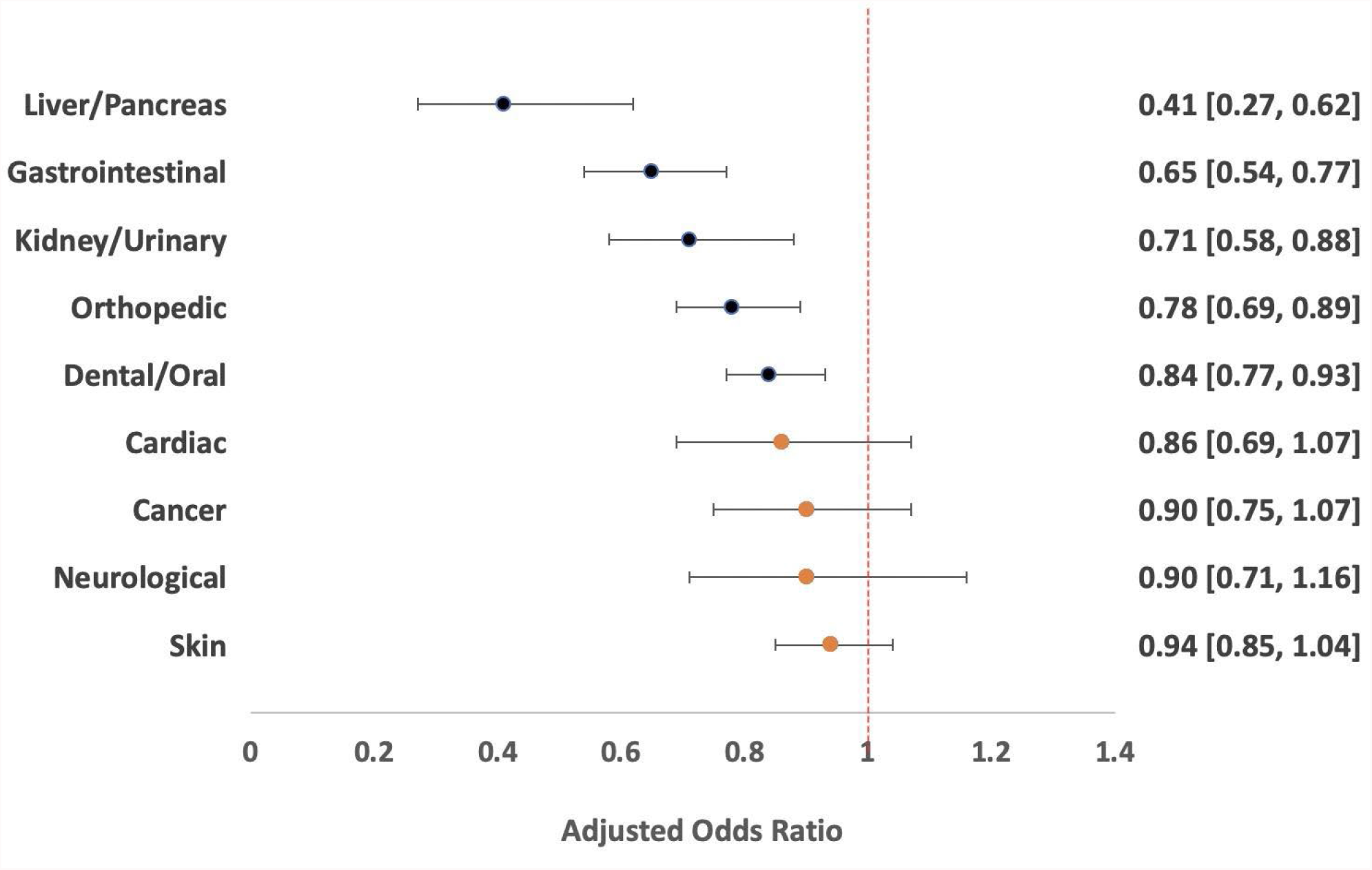
Summary of Results for Analysis of Health Conditions. Circles represent point estimates of adjusted odds ratios, with filled circles indicating statistically significant results. Bars represent 95% confidence intervals. Odds ratios less than 1 indicate lower odds of the outcome among dogs fed once daily; odds ratios greater than 1 indicate higher odds of the outcome among dogs fed once daily.

**Table 2.** Estimated odds ratios of specific health condition for dogs fed once per day compared to more frequently, adjusted for sex, age, breed for purebred dogs, and body size (as captured by weight) for mixed breed dogs. All analyses except Liver/Pancreas and Dental/Oral are also adjusted for omega-3 (or other fatty acid) supplementation.

| Health condition | Adjusted odds ratio | 95% CI | *p* |
| --- | --- | --- | --- |
| Liver/Pancreas | 0.41 | 0.27 - 0.61 | <0.001 |
| Gastrointestinal | 0.65 | 0.54 - 0.77 | <0.001 |
| Kidney/Urinary | 0.71 | 0.58 - 0.88 | 0.0012 |
| Orthopedic | 0.78 | 0.69 - 0.88 | <0.001 |
| Dental/Oral | 0.84 | 0.77 - 0.92 | <0.001 |
| Cardiac | 0.86 | 0.70 - 1.07 | 0.18 |
| Cancer | 0.90 | 0.75 - 1.07 | 0.24 |
| Neurological | 0.90 | 0.71 - 1.16 | 0.42 |
| Skin | 0.94 | 0.85 - 1.04 | 0.22 |

## Discussion

Using observational data from the Dog Aging Project, this is the largest study to date of feeding frequency conducted in companion dogs. We found that adult dogs fed once daily have better average cognitive scores and are less likely to have gastrointestinal, dental/oral, orthopedic, kidney/urinary, and liver/pancreas health conditions than dogs fed more frequently. While it is important to note that this study does not demonstrate causality, our observations are consistent with prior work in laboratory mice and observational studies in humans [61] suggesting that diets that restrict the timing of feeding are associated with better cognitive function and physical health.

In addition to being able to observe the animals in a naturalistic versus laboratory setting, one of the major strengths of our investigation is the large sample size of dogs included (CSLB assessment: 10,474 dogs; all other health conditions: 24,238 dogs). Furthermore, our statistical methods used flexible adjustment of continuous covariates (age, weight, and physical activity), reducing the possibility of residual confounding by these factors.

The key limitation of this work is that it is a cross-sectional analysis. Thus, we cannot rule out the possibility that dog owners shifted to more frequent feeding in response to health conditions, and observed associations are due in whole or part to reverse causality. This is a particular concern for gastrointestinal conditions and liver conditions, which are the two health conditions with the strongest observed associations. Such a shift would not be captured in the current dataset as owners reported on their dog’s current feeding frequency but did not provide information on feeding frequency history. In the future, we will gather this information through annual ‘snapshots’ since participants complete HLES each year. As the Dog Aging Project accrues longitudinal data over the next several years, investigations can compare dogs with different feeding frequencies who do not have a health condition and prospectively examine incidence of the condition. Such analyses can provide stronger evidence for a causal effect of feeding frequency on health.

It is plausible that once-daily feeding tends to result in lower overall caloric intake compared to more frequent feeding. Since we do not have data on caloric intake, we cannot analyze whether observed associations are mediated by caloric intake or reflect a possible effect of feeding frequency through other pathways. Caloric restriction has been previously reported to extend lifespan and improve health in Labrador retrievers maintained in a laboratory setting [5], although it has not yet been studied in other breeds of dog or in companion dogs.

This study has other limitations. All data are owner-reported and thus subject to error in recall and interpretation. However, while a given owner’s responses on their dog’s cognitive function, physical activity, and other health conditions might reflect individual differences in interpretation and reporting errors, it is unlikely that these would generate the specific associations we observed. We were also unable to account for dogs reported as fed once-daily but who received snacks and treats throughout the day. Although HLES gathers data on frequency of treats, we did not use these data because the caloric content of treats was unknown. Finally, due to the rarity of intact dogs in our sample, analyses included only spayed and neutered dogs. While age at spay or neuter might be an important factor for some health outcomes [62, 63], this information was not incorporated into our analyses because data on the timing of gonadectomy were not available with sufficient detail or completeness.

Studies of obesity, including possible associations with feeding frequency [23], will be an important area of future research. This investigation did not consider obesity because information on dogs’ body condition scores was not available. We anticipate that these data will be available in the future when owners share their dogs’ veterinary electronic medical records (VEMR) with the Dog Aging Project.

Given the limitations of this cross-sectional, observational study, the results of this investigation should not be used to make decisions about the feeding or clinical care of companion dogs.

However, if supported by future studies, it may be prudent to revisit the currently predominant recommendation that adult dogs be fed twice daily. The rationale for twice-daily feeding in dogs is obscure (although see [64]), and our study suggests that more frequent feeding may, in fact, be suboptimal for several age-related health outcomes.

We view these results as an exciting first step of an ongoing exploration of the impact of diet on companion dogs living in human environments. Given the intense interest in, and popularization of, “longevity diets” such as intermittent fasting and time-restricted feeding, these types of studies in dogs are both timely and important. We believe these studies will ultimately offer insights into factors that promote health and longevity for both dogs and humans.

## Author Contributions

All authors contributed to writing – review & editing. E.B.: conceptualization, methodology, data curation, writing – original draft, and project administration. Z.Z.: conceptualization, methodology, formal analysis, and visualization. K.T.: conceptualization and data curation. B.M.: data curation. DAP consortium: resources. M.K.: conceptualization, writing – original draft, and funding acquisition. K.K.: conceptualization, methodology, formal analysis, data curation, writing – original draft, visualization, project administration, and supervision.

## Dog Aging Project Consortium Authors

Joshua M. Akey, Brooke Benton, Elhanan Borenstein, Marta G. Castelhano, Amanda E. Coleman, Kate E. Creevy, Kyle Crowder, Matthew D. Dunbar, Virginia R. Fajt, Annette L. Fitzpatrick, Erica C. Jonlin, Unity Jeffrey, Elinor K. Karlsson, Jonathan M. Levine, Jing Ma, Robyn McClelland, Daniel E.L. Promislow, Audrey Ruple, Stephen M. Schwartz, Sandi Shrager, Noah Snyder-Mackler, Silvan R. Urfer, Benjamin S. Wilfond

## Acknowledgments

The Dog Aging Project thanks study participants, their dogs, and community veterinarians for their important contributions. The Dog Aging Project is supported by U19AG057377 and R24AG073137 from the National Institute on Aging, a part of the National Institutes of Health, and by additional grants and private donations. The content is solely the responsibility of the authors and does not necessarily represent the official views of the National Institutes of Health.

## Data availability statement

These data are housed on the Terra platform at the Broad Institute of MIT and Harvard.

## Code availability statement

This study did not use custom code or mathematical algorithms.

## Supplementary Information captions

Supplementary Information 1: Summary of the inclusion and exclusion criteria for subjects across all analyses, including guidelines for how all relevant variables were coded.

Supplementary Information 2: Criteria for determining whether or not a dog had a history of training (coded as a binary variable).

Supplementary Information 3: Regression outputs from the CSLB analysis, as well as the regression outputs from the health outcome analyses (both primary and secondary).

Supplementary Information 4: Complete list of purebred dogs (*n* = 76) included in the CSLB analysis, with sample sizes.

Supplementary Information 5: Complete list of purebred dogs (*n* = 100) included in analysis of health conditions, with sample sizes.

