## Supplementary Information 1 for "Once-daily feeding is associated with better health in companion dogs: Results from the Dog Aging Project"

**Logic behind inclusion/exclusion criteria**

| **Subset** | **Variable type** | **Measure** | **Rule** | **Justification** |
| --- | --- | --- | --- | --- |
| *All analyses* | Exposure (feeding frequency) | HLES: Times per day being fed | Analyze as a binary variable: once per day vs. more frequently (combining the twice, three or more, & free fed categories) | Exposure of interest |
|  | Screening | HLES: Consistency of daily diet | Exclude dogs with diets reported as "not at all consistent" | Reported feeding frequency (exposure) needs to be consistent to look for associations with outcomes |
|  | Screening | Sex, Spayed or Neutered | Exclude un-Neutered and un-Spayed dogs | Very small numbers (95% of overall DAP sample consists of altered dogs) |
|  | Screening | Age | Exclude dogs >= 18 years old | Difficult to verify; outliers |
|  | Precision variable | Breed for purebred dogs; Weight for mixed breed dogs | Purebred dogs: analyze breed as a categorical variable, including only breeds with more than 10 subjects total and with at least one affected and one unaffected dog; Mixed breed dogs: analyze weight as a continuous variable | Previous work has shown that, for certain conditions, prevalence is much higher in certain breeds as compared to others; also, a proxy for body size |
|  | Precision variable | Sex | Analyze as a binary variable: male, female | Can affect almost any outcome |
| *Cognitive analysis only* | Outcome (cognitive) | CSLB score | Analyze as a continuous variable | Owners must have completed the CSLB survey (in addition to HLES) |
|  | Potential confounder | Age | Exclude dogs < 6 years old; Analyze as continuous variable | We don't expect to see meaningful declines in cognitive function prior to 6 years of age |
| *Cognitive analysis only continued* | Precision variable | HLES: Primary activity | Analyze as a binary variable: companion/pet vs. history of training (i.e., participation in an activity that involves training/cognitive enrichment); exclude dogs where this categorization cannot be determined (see SI_2 for details) | Included as a proxy for training/cognitive enrichment, which has been hypothesized to affect cognition in dogs |
|  | Potential confounder | HLES: Physical activity | Analyze as a principal component score of 3 relevant measures: lifestyle activity level, average activity intensity level, and average daily time spent physically active | Previous work has shown that physical activity level has positive effects on cognitive outcomes |
|  | Potential confounder | Omega 3 or other fatty acids, given as a separate supplement at least daily | Analyze as a binary variable (affected/unaffected) | Previous work has shown that omega-3 (i.e., fish oil) has positive effects on cognitive outcomes |
| *Health analyses only* | Outcome (health) | Skin disorders | Analyze as a binary variable (affected/unaffected); Exclude disorders related to situational factors | Disorders linked to transient situational factors, including fleas and ticks, are excluded |
|  | Outcome (health) | Dental or oral disease | Analyze as a binary variable (affected/unaffected); Exclude congenital disorders and those related to situational factors | Congenital disorders are excluded as they appear prior to the exposure (feeding frequency); disorders linked to transient situational factors, including infectious diseases and trauma, are excluded |
| *Health analyses only continued* | Outcome (health) | Orthopedic disorders | Analyze as a binary variable (affected/unaffected); Exclude congenital disorders and those related to situational factors | Congenital disorders are excluded as they appear prior to the exposure (feeding frequency); disorders linked to transient situational factors, including infectious diseases and trauma, are excluded |
|  | Outcome (health) | Gastrointestinal disorders | Analyze as a binary variable (affected/unaffected); Exclude congenital disorders and those related to situational factors | Congenital disorders are excluded as they appear prior to the exposure (feeding frequency); disorders linked to transient situational factors, including infectious diseases and trauma, are excluded |
|  | Outcome (health) | Kidney or urinary disorders | Analyze as a binary variable (affected/unaffected); Exclude congenital disorders | Congenital disorders are excluded as they appear prior to the exposure (feeding frequency) |
|  | Outcome (health) | Cancer or tumors | Analyze as a binary variable (affected/unaffected | Health outcome conceivably related to feeding frequency |
|  | Outcome (health) | Cardiac disorders | Analyze as a binary variable (affected/unaffected); Exclude congenital disorders and those related to situational factors | Congenital disorders are excluded as they appear prior to the exposure (feeding frequency); disorders linked to transient situational factors, including infectious diseases and trauma, are excluded |
| *Health analyses only continued* | Outcome (health) | Neurological disorders | Analyze as a binary variable (affected/unaffected); Exclude disorders related to situational factors | Disorders linked to transient situational factors, including infectious diseases and trauma, are excluded |
|  | Outcome (health) | Liver or pancreas disorders | Analyze as a binary variable (affected/unaffected); Exclude congenital disorders | Congenital disorders are excluded as they appear prior to the exposure (feeding frequency) |
|  | Screening outcomes | HLES: Dogs who have been diagnosed with condition(s) | Exclude Trauma category as an outcome | Situational / not a disease outcome |
|  | Screening outcomes | HLES: Dogs who have been diagnosed with condition(s) | Exclude Toxic and controlled substance category as an outcome | Situational / not a disease outcome |
|  | Screening outcomes | HLES: Dogs who have been diagnosed with condition(s) | Exclude Infectious or parasitic disease category as an outcome | Situational / based on environmental factors (e.g. exposure to tick/virus/bacteria) |
|  | Screening outcomes | HLES: Dogs who have been diagnosed with condition(s) | Exclude Endocrine disorders category as an outcome | For the biggest subset (hypothyroidism), there isn’t a compelling case as to why feeding frequency would be important. Other categories that we would be interested in (e.g., Cushing’s disease, diabetes) don’t have a big enough sample size |
| *Health analyses only continued* | Screening outcomes | HLES: Dogs who have been diagnosed with condition(s) | Exclude Respiratory disorders category as an outcome | Sample size is too small |
|  | Screening outcomes | HLES: Dogs who have been diagnosed with condition(s) | Exclude Immune-mediated disorders category as an outcome | Sample size is too small |
|  | Screening outcomes | HLES: Dogs who have been diagnosed with condition(s) | Exclude Hematopoietic disorders category as an outcome | Sample size is too small |
|  | Screening outcomes | HLES: Dogs who have been diagnosed with condition(s) | Exclude Reproductive system disorders category as an outcome | Sample size is too small |
|  | Screening outcomes | HLES: Dogs who have been diagnosed with condition(s) | Exclude Eye disorders category as an outcome | Unlikely to be affected by feeding frequency |
| *Health analyses only continued* | Screening outcomes | HLES: Dogs who have been diagnosed with condition(s) | Exclude Ear, nose, and throat disorders as an outcome | Unlikely to be affected by feeding frequency |
|  | Potential confounder | Age | Exclude dogs < 1 year old | Feeding frequency differs for puppies as compared to adults |
| *Subset of health analyses only: Skin, Orthopedic, Gastrointestinal, Kidney/Urinary, Cancer, Cardiac, Neurological* | Potential confounder | Omega-3 or other fatty acids, given as a separate supplement at least daily | Analyze as a binary variable (affected/unaffected) | Previous work has shown that omega-3 (i.e., fish oil) has positive effects on outcomes in these health areas |

**Dental/Oral: Guidelines for coding of affected/unaffected**

| **Dental/Oral Category** | **Designation (affected or unaffected)** | **Justification** |
| --- | --- | --- |
| Dental calculus | Affected | Health outcome conceivably related to feeding frequency |
| Extracted teeth | Affected | Health outcome conceivably related to feeding frequency |
| Fractured teeth | Affected | Health outcome conceivably related to feeding frequency |
| Gingivitis (red, puffy gums) | Affected | Health outcome conceivably related to feeding frequency |
| Underbite | Not affected | Congenital |
| Other | Varies | See below |
| Retained deciduous (baby) teeth | Not affected | Congenital |
| Overbite | Not affected | Congenital |
| Oronasal fistula | Not affected | Congenital |
| Sialocele | Not affected | Caused by trauma |
| Masticatory myositis | Not affected | Health outcome not plausibly related to feeding frequency |

**Guidelines for write-in answers from dental/oral ‘other’ category**

| **Dental/Oral ‘Other’ Category** | **Designation (affected or unaffected)** | **Justification** |
| --- | --- | --- |
| Anatomic abnormalities (e.g., malocclusion, elongated palate) | Not affected | Congenital |
| Trauma (e.g., fractures, extracted teeth) | Not affected | Situational |
| Congenital infections | Not affected | Congenital |

**Skin: Guidelines for coding of affected/unaffected**

| **Skin Category** | **Designation (affected or unaffected)** | **Justification** |
| --- | --- | --- |
| Seasonal allergies | Affected | Health outcome conceivably related to feeding frequency |
| Pruritis (itchy skin) | Affected | Health outcome conceivably related to feeding frequency |
| Sebaceous cysts | Affected | Health outcome conceivably related to feeding frequency |
| Food/med allergies that affect the skin | Affected | Health outcome conceivably related to feeding frequency |
| Chronic or recurrent hot spots | Affected | Health outcome conceivably related to feeding frequency |
| Fleas | Not affected | Situational |
| Other | Varies | See below |
| Atopic dermatitis (atopy) | Affected | Health outcome conceivably related to feeding frequency |
| Ticks | Not affected | Situational |
| Chronic or recurrent skin infections | Affected | Health outcome conceivably related to feeding frequency; included even though infectious because it's persistent/chronic/recurring |
| Flea allergy dermatitis | Not affected | Situational |
| Contact dermatitis | Affected | Health outcome conceivably related to feeding frequency |
| Alopecia (hair loss) | Affected | Health outcome conceivably related to feeding frequency |
| Non-specific dermatitis | Affected | Health outcome conceivably related to feeding frequency |
| Lick granuloma | Affected | Health outcome conceivably related to feeding frequency |
| Pyoderma or bacterial dermatitis | Affected | Health outcome conceivably related to feeding frequency |
| Systemic demodectic mange | Affected | Health outcome conceivably related to feeding frequency |
| Sarcoptic mange | Not affected | Parasitic |
| Seborrhea (greasy skin) | Affected | Health outcome conceivably related to feeding frequency |
| Pododermatitis | Affected | Health outcome conceivably related to feeding frequency |
| Discoid lupus erythematosus (DLE) | Affected | Health outcome conceivably related to feeding frequency |
| Ichthyosis | Affected | Health outcome conceivably related to feeding frequency |
| Sebaceous adenitis | Affected | Health outcome conceivably related to feeding frequency |
| Pemphigus foliaceus (PF) | Affected | Health outcome conceivably related to feeding frequency |
| Systemic lupus erythematosus (SLE) | Affected | Health outcome conceivably related to feeding frequency |
| Panepidermal pustular pemphigus (PPP) | Affected | Health outcome conceivably related to feeding frequency |
| Pemphigus erythematosus (PE) | Affected | Health outcome conceivably related to feeding frequency |
| Pemphigus vulgaris (PV) | Affected | Health outcome conceivably related to feeding frequency |

**Guidelines for write-in answers from skin ‘other’ category:**

| **Skin ‘Other’ Category** | **Designation (affected or unaffected)** | **Justification** |
| --- | --- | --- |
| Staph infections | Not affected | Infectious disease |
| Ringworm infections | Not affected | Infectious disease |
| Scarring | Not affected | Situational |
| Infections (e.g., yeast, skin) | Not affected | Infectious disease |
| Lice | Not affected | Parasitic |
| Mites | Not affected | Parasitic |
| Bot fly | Not affected | Parasitic |
| Sunburn / sun allergy | Not affected | Situational |
| Bee sting allergy | Not affected | Situational |
| Reactions secondary to drugs or vaccines | Not affected | Situational |

**Orthopedic: Guidelines for coding of affected/unaffected**

| **Orthopedic Category** | **Designation (affected or unaffected)** | **Justification** |
| --- | --- | --- |
| Osteoarthritis | Affected | Health outcome conceivably related to feeding frequency |
| Cruciate ligament rupture | Affected | Health outcome conceivably related to feeding frequency |
| Patellar luxation | Affected | Health outcome conceivably related to feeding frequency |
| Hip dysplasia | Affected | Health outcome conceivably related to feeding frequency |
| Other | Varies | See below |
| Lameness (chronic or recurrent) | Affected | Health outcome conceivably related to feeding frequency |
| Degenerative joint disease | Affected | Health outcome conceivably related to feeding frequency |
| Intervertebral disc disease (IVDD) | Affected | Health outcome conceivably related to feeding frequency |
| Elbow dysplasia | Affected | Health outcome conceivably related to feeding frequency |
| Rheumatoid arthritis | Affected | Health outcome conceivably related to feeding frequency |
| Spondylosis | Affected | Health outcome conceivably related to feeding frequency |
| Growth deformity | Affected | Health outcome conceivably related to feeding frequency |
| Panosteitis | Affected | Health outcome conceivably related to feeding frequency |
| Osteochondritis dissecans (OCD) | Affected | Health outcome conceivably related to feeding frequency |
| Carpal subluxation syndrome | Affected | Health outcome conceivably related to feeding frequency |
| Dwarfism | Not affected | Congenital |
| Osteomyelitis | Not affected | Infectious disease |

**Guidelines for write-in answers from orthopedic ‘other’ category:**

| **Orthopedic ‘Other’ Category** | **Designation (affected or unaffected)** | **Justification** |
| --- | --- | --- |
| Trauma (e.g., limb removal) | Not affected | Situational |
| Congenital (e.g., missing limbs/deformities at birth) | Not affected | Congenital |
| Congenital dysplasia | Not affected | Congenital (if congenital is not specified, we designated dysplasia as affected) |

**Gastrointestinal: Guidelines for coding of affected/unaffected**

| **Gastrointestinal Category** | **Designation (affected or unaffected)** | **Justification** |
| --- | --- | --- |
| Chronic or recurrent diarrhea | Affected | Health outcome conceivably related to feeding frequency |
| Anal sac impaction | Affected | Health outcome conceivably related to feeding frequency |
| Foreign body ingestion or blockage | Not affected | Situational |
| Food or medicine allergies | Affected | Health outcome conceivably related to feeding frequency |
| Other | Varies | See below |
| Irritable bowel syndrome (IBS) | Affected | Health outcome conceivably related to feeding frequency |
| Chronic or recurrent vomiting | Affected | Health outcome conceivably related to feeding frequency |
| Hemorrhagic gastroenteritis (HGE) | Affected | Health outcome conceivably related to feeding frequency |
| Other allergies | Affected | Health outcome conceivably related to feeding frequency |
| Bilious vomiting syndrome | Affected | Health outcome conceivably related to feeding frequency |
| Fecal incontinence | Affected | Health outcome conceivably related to feeding frequency |
| Constipation | Affected | Health outcome conceivably related to feeding frequency |
| Bloat with torsion (GDV) | Affected | Health outcome conceivably related to feeding frequency |
| Idiopathic canine colitis | Affected | Health outcome conceivably related to feeding frequency |
| Megaesophagus | Affected | Health outcome conceivably related to feeding frequency |
| Protein-losing enteropathy (PLE) | Affected | Health outcome conceivably related to feeding frequency |
| Malabsorptive disorder | Affected | Health outcome conceivably related to feeding frequency |
| Lymphangiectasia | Affected | Health outcome conceivably related to feeding frequency |
| Pyloric stenosis | Not affected | Congenital |

**Guidelines for write-in answers from gastrointestinal ‘other’ category:**

| **Gastrointestinal ‘Other’ Category** | **Designation (affected or unaffected)** | **Justification** |
| --- | --- | --- |
| Infectious diseases | Not affected | Infectious |
| Trauma | Not affected | Situational |

**Cancer: Guidelines for coding of affected/unaffected**

| **Cancer Category** | **Designation (affected or unaffected)** | **Justification** |
| --- | --- | --- |
| Don't know | Affected | Health outcome conceivably related to feeding frequency |
| Mast cell tumor | Affected | Health outcome conceivably related to feeding frequency |
| Lipoma | Affected | Health outcome conceivably related to feeding frequency |
| Other | Affected | Health outcome conceivably related to feeding frequency |
| Soft tissue sarcoma | Affected | Health outcome conceivably related to feeding frequency |
| Melanoma | Affected | Health outcome conceivably related to feeding frequency |
| Carcinoma | Affected | Health outcome conceivably related to feeding frequency |
| Lymphoma/lymphosarcoma | Affected | Health outcome conceivably related to feeding frequency |
| Sarcoma | Affected | Health outcome conceivably related to feeding frequency |
| Adenoma | Affected | Health outcome conceivably related to feeding frequency |
| Adenocarcinoma | Affected | Health outcome conceivably related to feeding frequency |
| Hemangiosarcoma | Affected | Health outcome conceivably related to feeding frequency |
| Histiocytoma | Affected | Health outcome conceivably related to feeding frequency |
| Basal cell tumor | Affected | Health outcome conceivably related to feeding frequency |
| Osteosarcoma | Affected | Health outcome conceivably related to feeding frequency |
| Epidermoid cyst | Affected | Health outcome conceivably related to feeding frequency |
| Hemangioma | Affected | Health outcome conceivably related to feeding frequency |
| Squamous cell carcinoma | Affected | Health outcome conceivably related to feeding frequency |
| Papilloma | Affected | Health outcome conceivably related to feeding frequency |
| Sebaceous adenoma | Affected | Health outcome conceivably related to feeding frequency |
| Fibrosarcoma | Affected | Health outcome conceivably related to feeding frequency |
| Plasmacytoma | Affected | Health outcome conceivably related to feeding frequency |
| Epulides | Affected | Health outcome conceivably related to feeding frequency |
| Peripheral nerve sheath tumor | Affected | Health outcome conceivably related to feeding frequency |
| Transitional cell carcinoma | Affected | Health outcome conceivably related to feeding frequency |
| Leukemia | Affected | Health outcome conceivably related to feeding frequency |
| Histiocytic sarcoma | Affected | Health outcome conceivably related to feeding frequency |
| Insulinoma | Affected | Health outcome conceivably related to feeding frequency |
| Chondrosarcoma | Affected | Health outcome conceivably related to feeding frequency |
| Cystadenoma | Affected | Health outcome conceivably related to feeding frequency |
| Leiomyoma | Affected | Health outcome conceivably related to feeding frequency |
| Meningioma | Affected | Health outcome conceivably related to feeding frequency |
| Leiomyosarcoma | Affected | Health outcome conceivably related to feeding frequency |
| Multiple myeloma | Affected | Health outcome conceivably related to feeding frequency |
| Thymoma | Affected | Health outcome conceivably related to feeding frequency |

**Kidney/Urinary: Guidelines for coding of affected/unaffected**

| **Kidney/Urinary Category** | **Designation (affected or unaffected)** | **Justification** |
| --- | --- | --- |
| Urinary tract infection | Affected | Health outcome conceivably related to feeding frequency |
| Urinary incontinence | Affected | Health outcome conceivably related to feeding frequency |
| Urinary crystals or stones in bladder | Affected | Health outcome conceivably related to feeding frequency |
| Chronic kidney disease | Affected | Health outcome conceivably related to feeding frequency |
| Other | Varies | See below |
| Acute kidney failure | Affected | Health outcome conceivably related to feeding frequency |
| Proteinuria | Affected | Health outcome conceivably related to feeding frequency |
| Kidney stones | Affected | Health outcome conceivably related to feeding frequency |
| Pyelonephritis (kidney infection) | Affected | Health outcome conceivably related to feeding frequency |
| Renal dysplasia | Not affected | Congenital |
| Ectopic ureter | Not affected | Congenital |
| Bladder prolapse | Affected | Health outcome conceivably related to feeding frequency |
| Urethral prolapse | Affected | Health outcome conceivably related to feeding frequency |
| Tubular disorder | Affected | Health outcome conceivably related to feeding frequency |

**Guidelines for write-in answers from kidney/urinary ‘other’ category:**

| **Kidney/Urinary ‘Other’ Category** | **Designation (affected or unaffected)** | **Justification** |
| --- | --- | --- |
| Trauma (e.g., hematoma from injury) | Not affected | Situational |
| Acute toxic | Not affected | Situational |
| Behavioral (e.g., submissive peeing, excitable peeing) | Not affected | Categorized incorrectly |

**Cardiac: Guidelines for coding of affected/unaffected**

| **Cardiac Category** | **Designation (affected or unaffected)** | **Justification** |
| --- | --- | --- |
| Murmur | Affected | Health outcome conceivably related to feeding frequency |
| Valve disease | Affected | Health outcome conceivably related to feeding frequency |
| Congestive heart failure | Affected | Health outcome conceivably related to feeding frequency |
| Arrhythmia | Affected | Health outcome conceivably related to feeding frequency |
| Cardiomyopathy | Affected | Health outcome conceivably related to feeding frequency |
| Other | Varies | See below |
| Hypertension (high blood pressure) | Affected | Health outcome conceivably related to feeding frequency |
| Pulmonary hypertension | Affected | Health outcome conceivably related to feeding frequency |
| Subaortic stenosis | Not affected | Congenital |
| Pulmonic stenosis | Not affected | Congenital |
| Endocarditis | Not affected | Infectious disease |
| Pericardial effusion | Affected | Health outcome conceivably related to feeding frequency |

**Guidelines for write-in answers from cardiac ‘other’ category:**

| **Cardiac ‘Other’ Category** | **Designation (affected or unaffected)** | **Justification** |
| --- | --- | --- |
| Heartworm | Not affected | Infectious |
| Toxic (e.g., enlarged heart due to rat poison) | Not affected | Situational |
| Patent ductus arteriosus | Not affected | Congenital |

**Neurological: Guidelines for coding of affected/unaffected**

| **Neurological Category** | **Designation (affected or unaffected)** | **Justification** |
| --- | --- | --- |
| Cauda equina syndrome | Affected | Health outcome conceivably related to feeding frequency |
| Degenerative myelopathy | Affected | Health outcome conceivably related to feeding frequency |
| Dementia or senility | Affected | Health outcome conceivably related to feeding frequency |
| Discospondylitis | Not affected | Infectious disease |
| Fibrocartilaginous embolism (FCE) | Affected | Health outcome conceivably related to feeding frequency |
| Horner's syndrome | Affected | Health outcome conceivably related to feeding frequency |
| Intervertebral disc disease (IVDD) (Neurologic) | Affected | Health outcome conceivably related to feeding frequency |
| Laryngeal paralysis (Neurologic) | Affected | Health outcome conceivably related to feeding frequency |
| Limb paralysis | Affected | Health outcome conceivably related to feeding frequency |
| Myasthenia gravis | Affected | Health outcome conceivably related to feeding frequency |
| Polyneuropathy | Affected | Health outcome conceivably related to feeding frequency |
| Seizures (including epilepsy) | Affected | Health outcome conceivably related to feeding frequency |
| Vestibular disease | Affected | Health outcome conceivably related to feeding frequency (confirmed that we are including all types, as HLES distinguishes central, peripheral, and other) |
| Wobbler syndrome | Affected | Health outcome conceivably related to feeding frequency |
| Other | Varies | See below |

**Guidelines for write-in answers from neurological ‘other’ category:**

| **Neurological ‘Other’ Category** | **Designation (affected or unaffected)** | **Justification** |
| --- | --- | --- |
| Exercise induced collapse | Not affected | Situational |
| Toxic (e.g., reaction to flea/tick preventative or vaccine, resulting from ingestion of fungus) | Not affected | Situational |
| Trauma (e.g., myelopathy) | Not affected | Situational |

**Liver/Pancreas: Guidelines for coding of affected/unaffected**

| **Live/Pancreas Category** | **Designation (affected or unaffected)** | **Justification** |
| --- | --- | --- |
| Pancreatitis | Affected | Health outcome conceivably related to feeding frequency |
| Other | Varies | See below |
| Chronic inflammatory liver disease | Affected | Health outcome conceivably related to feeding frequency |
| Exocrine pancreatic insufficiency (EPI) | Affected | Health outcome conceivably related to feeding frequency |
| Gall bladder mucocele | Affected | Health outcome conceivably related to feeding frequency |
| Gall bladder surgery | Affected | Health outcome conceivably related to feeding frequency |
| Microvascular dysplasia | Not affected | Congenital |
| Biliary obstruction | Affected | Health outcome conceivably related to feeding frequency |
| Gall bladder rupture | Affected | Health outcome conceivably related to feeding frequency |
| Portosystemic shunt | Not affected | Congenital |

**Guidelines for write-in answers from liver/pancreas ‘other’ category:**

| **Liver/Pancreas ‘Other’ Category** | **Designation (affected or unaffected)** | **Justification** |
| --- | --- | --- |
| Liver infections | Not affected | Infectious |
| Gall bladder infection | Not affected | Infectious |
| Portal vein hypoplasia | Not affected | Congenital |
| Trauma or drug induced (e.g., phenobarbital-induced liver enzyme elevations; liver damage by doxycycline) | Not affected | Situational |
