## Supplementary Information 2 for "Once-daily feeding is associated with better health in companion dogs: Results from the Dog Aging Project"

Below we provide the detailed criteria for whether a dog had a history of training (coded as a binary variable), determined by considering HLES responses for a dog’s ‘Primary activity’ and ‘Secondary activity’.

**If only one activity was reported:**

| **Category** | **History of training (1/0)** | **Justification** |
| --- | --- | --- |
| Companion animal or pet | 0 | No explicit training required |
| Service dog (Seeing eye dog, Hearing or signal dog, Wheelchair service dog) | 1 | Requires training |
| Assistance or therapy dog | Varies | See subcategories below |
| Obedience | 1 | Requires training |
| Agility | 1 | Requires training |
| Working (herding, guarding, etc.) | 1 | Requires training |
| Hunting | 1 | Requires training |
| Show | 1 | Requires training |
| Search & Rescue | 1 | Requires training |
| Field Trials | 1 | Requires training |
| Breeding | 0 | No explicit training required |
| Other | Varies | See guidelines below |

**Assistance or therapy dog subcategories:**

| **Category** | **History of training (1/0)** | **Justification** |
| --- | --- | --- |
| Community therapy dog | 1 | Requires training |
| Emotional support dog | 0 | No explicit training required |

**Guidelines for write-in answers from ‘other’ categories (including Other, Other medical service dog, Other health assistance dog):**

| **Category** | **History of training (1/0)** | **Justification** |
| --- | --- | --- |
| Not enough information (e.g. demonstration dog but no indication of for what) | Excluded | Unclear if training is involved |
| Athlete | Excluded | Unclear if training is involved or just recreational |
| Mobility support | 1 | Requires training |
| PTSD service dog | 1 | Requires training |
| Facility dog (e.g., courtroom dog) | 1 | Requires training |
| Psychiatric service dog | 1 | Requires training |
| Medical alert dog (e.g., diabetic alert) | 1 | Requires training |
| Nose work/scent work | 1 | Requires training |
| Police K9 | 1 | Requires training |
| Sled dog mushing | 1 | Requires training |
| Mentions training or obedience work | 1 | Requires training |
| Autism service dog | 1 | Requires training |
| Greyhound/track racing | 1 | Requires training |
| Flyball | 1 | Requires training |
| Dock diving | 1 | Requires training |
| Disc | 1 | Requires training |
| Rally | 1 | Requires training |
| Barn Hunt | 1 | Requires training |
| Frisbee | 1 | Requires training |
| Reading therapy | 0 | Unless specifically designated as a 'therapy dog' or as a participant in a recognized therapy dog program; otherwise: no explicit training required |
| Visits to hospitals/schools/nursing homes | 0 | Unless specifically designated as a 'therapy dog'; otherwise: no explicit training required |
| Blood donor | 0 | No explicit training required |
| Exercise/hiking/joring/walking buddy | 0 | No explicit training required |
| Whimsical, tongue-in-cheek responses (e.g., keeps me smiling) | 0 | No explicit training required |
| Office dog/greeter | 0 | No explicit training required |
| Guard dog/Watch dog | 0 | No explicit training required |
| Unofficial/"Untrained" therapy dog | 0 | No explicit training required |
| Comfort dog / Mental Support | 0 | No explicit training required |
| Distraction from pain | 0 | No explicit training required |
| Lure Coursing | 0 | No explicit training required |

**Guidelines for dogs with both a primary and secondary activity:**

- If at least one activity was determined to involve training, the dog was coded as having a history of training (1)
- If both activities were determined to not need explicit training, the dog was coded as *not* having a history of training (0)
- If one activity was determined to not need explicit training and the other was unable to be determined, we were unable to determine the ‘history of training’ score and these dogs were excluded from the CSLB analysis
