## Supplementary Information 3 for "Once-daily feeding is associated with better health in companion dogs: Results from the Dog Aging Project"

The "breed effect" in a fitted regression model is relative to a 50-lb mixed-breed dog. Due to flexible modeling using natural cubic splines for continuous variables, we do not report regression coefficients for continuous variables.

For health outcomes, the primary analysis used conditional logistic regression with the Efron approximation. Secondary analysis used ordinary logistic regression and data on mixed-breed dogs and dogs from the 10 most populous breeds in the dataset (75.5% of total analytic dataset).

**CSLB outcome: Linear regression results**

Primary CSLB Analysis: Linear regression, *n* = 10,474.

| **Explanatory variable** | **Estimated odds ratio** | **95% Confidence Interval** | | ***p*** | **Degrees of freedom** | |
| --- | --- | --- | --- | --- | --- | --- |
| feed once | -0.62 | (-0.97, -0.27) | <0.001 | | | 1 |
| sex: female | -0.11 | (-0.30, 0.080) | 0.26 | | | 1 |
| cognitive training | -0.38 | (-0.63, -0.14) | 0.0023 | | | 1 |
| fatty acid | -0.16 | (-0.39, 0.062) | 0.15 | | | 1 |
| physical activity level |  |  | <0.001 | | | 4 |
| age |  |  | <0.001 | | | 4 |
| weight |  |  | 0.37 | | | 4 |
| Airedale Terrier | -0.099 | (-1.81, 1.61) | 0.91 | | | 1 |
| Alaskan Malamute | 1.47 | (-1.32, 4.26) | 0.30 | | | 1 |
| American Eskimo Dog | 0.40 | (-2.27, 3.07) | 0.77 | | | 1 |
| American Pitbull Terrier | -1.19 | (-2.62, 0.25) | 0.11 | | | 1 |
| American Staffordshire Terrier | -1.50 | (-3.61, 0.61) | 0.16 | | | 1 |
| Australian Cattle Dog | 0.80 | (-0.63, 2.24) | 0.27 | | | 1 |
| Australian Shepherd | 0.30 | (-0.51, 1.10) | 0.47 | | | 1 |
| Basset Hound | 0.41 | (-0.99, 1.82) | 0.56 | | | 1 |
| Beagle | -0.14 | (-1.09, 0.82) | 0.78 | | | 1 |
| Bernese Mountain Dog | -0.86 | (-2.23, 0.50) | 0.22 | | | 1 |
| Bichon Frise | 0.16 | (-1.58, 1.90) | 0.86 | | | 1 |
| Border Collie | -0.53 | (-1.45, 0.39) | 0.26 | | | 1 |
| Border Terrier | 0.10 | (-1.32, 1.52) | 0.89 | | | 1 |
| Boston Terrier | 1.43 | (-0.36, 3.21) | 0.12 | | | 1 |
| Boxer | 0.30 | (-0.65, 1.25) | 0.54 | | | 1 |
| Brittany | -0.12 | (-1.61, 1.36) | 0.87 | | | 1 |
| Bull Terrier | 0.10 | (-1.62, 1.82) | 0.91 | | | 1 |
| Bull Dog | 1.61 | (-0.59, 3.81) | 0.15 | | | 1 |
| Cairn Terrier | -0.46 | (-2.03, 1.11) | 0.57 | | | 1 |
| Carolina Dog | 0.76 | (-1.18, 2.70) | 0.44 | | | 1 |
| Catahoula Leopard Dog | -3.37 | (-6.38, -0.36) | 0.028 | | | 1 |
| Cavalier King Charles Spaniel | -0.47 | (-1.55, 0.62) | 0.40 | | | 1 |
| Chesapeake Bay Retriever | -1.40 | (-3.18, 0.37) | 0.12 | | | 1 |
| Chihuahua | -2.70 | (-3.79, -1.61) | <0.001 | | | 1 |
| Cocker Spaniel | 0.31 | (-1.19, 1.82) | 0.68 | | | 1 |
| Collie | -0.92 | (-2.92, 1.07) | 0.37 | | | 1 |
| Dachshund | -1.00 | (-1.81, -0.19) | 0.016 | | | 1 |
| Dalmatian | -1.15 | (-3.62, 1.31) | 0.36 | | | 1 |
| Doberman Pinscher | 0.20 | (-0.53, 0.94) | 0.59 | | | 1 |
| English Cocker Spaniel | 1.18 | (-3.03, 5.39) | 0.58 | | | 1 |
| English Setter | -0.36 | (-2.16, 1.45) | 0.70 | | | 1 |
| English Springer Spaniel | 1.26 | (-0.14, 2.66) | 0.078 | | | 1 |
| French Bulldog | 1.05 | (-0.24, 2.35) | 0.11 | | | 1 |
| German Shepherd | -0.43 | (-1.06, 0.20) | 0.18 | | | 1 |
| German Shorthaired Pointer | 0.45 | (-0.83, 1.73) | 0.49 | | | 1 |
| Golden Retriever | -0.77 | (-1.19, -0.35) | <0.001 | | | 1 |
| Great Dane | 1.14 | (-0.48, 2.75) | 0.17 | | | 1 |
| Great Pyrenees | 0.16 | (-1.23, 1.56) | 0.82 | | | 1 |
| Greyhound | -1.05 | (-2.15, 0.045) | 0.060 | | | 1 |
| Havanese | -0.62 | (-1.43, 0.19) | 0.13 | | | 1 |
| Italian Greyhound | -0.33 | (-1.93, 1.27) | 0.69 | | | 1 |
| Jack Russell Terrier | -0.19 | (-1.31, 0.94) | 0.74 | | | 1 |
| Keeshond | 0.61 | (-1.34, 2.55) | 0.54 | | | 1 |
| Labrador Retriever | -0.31 | (-0.73, 0.098) | 0.16 | | | 1 |
| Lhasa Apso | -2.41 | (-3.90, -0.96) | 0.0014 | | | 1 |
| Maltese | -0.52 | (-2.30, 1.26) | 0.57 | | | 1 |
| Miniature Schnauzer | 0.26 | (-1.10, 1.63) | 0.70 | | | 1 |
| Miniature American Shepherd | 0.36 | (-0.86, 1.58) | 0.56 | | | 1 |
| Miniature Pinscher | 0.45 | (-1.36, 2.26) | 0.63 | | | 1 |
| Newfoundland | 0.57 | (-0.84, 1.98) | 0.43 | | | 1 |
| Papillon | -0.38 | (-1.70, 0.94) | 0.58 | | | 1 |
| Parson Russell Terrier | 0.43 | (-1.89, 2.75) | 0.71 | | | 1 |
| Pekingese | -0.16 | (-2.54, 2.22) | 0.89 | | | 1 |
| Pembroke Welsh Corgi | 0.39 | (-0.61, 1.38) | 0.45 | | | 1 |
| Pomeranian | 0.095 | (-1.87, 2.06) | 0.92 | | | 1 |
| Poodle (Standard)  (≥30 lbs) | -0.24 | (-1.04, 0.56) | 0.56 | | | 1 |
| Poodle (Toy) | 1.28 | (-0.84, 3.41) | 0.24 | | | 1 |
| Portuguese Water Dog | -0.68 | (-2.08, 0.72) | 0.34 | | | 1 |
| Pug | 0.36 | (-0.82, 1.55) | 0.55 | | | 1 |
| Rat Terrier | 0.93 | (-1.31, 3.18) | 0.42 | | | 1 |
| Rhodesian Ridgeback | -0.24 | (-0.97, 0.49) | 0.52 | | | 1 |
| Rottweiler | -0.52 | (-1.96, 0.93) | 0.49 | | | 1 |
| Samoyed | -0.57 | (-1.49, 0.34) | 0.22 | | | 1 |
| Scottish Terrier | -0.74 | (-3.28, 1.80) | 0.57 | | | 1 |
| Shetland Sheepdog | -0.71 | (-1.91, 0.49) | 0.25 | | | 1 |
| Shiba Inu | 3.05 | (0.72, 5.37) | 0.010 | | | 1 |
| Shih Tzu | 1.14 | (-0.46, 2.75) | 0.16 | | | 1 |
| Siberian Husky | 0.23 | (-1.26, 1.74) | 0.76 | | | 1 |
| Poodle (Standard)  (<30 lbs) | 0.33 | (-1.92, 2.58) | 0.77 | | | 1 |
| Soft Coated Wheaten Terrier | 2.35 | (0.95, 3.74) | <0.001 | | | 1 |
| Vizsla | 0.15 | (-0.96, 1.25) | 0.79 | | | 1 |
| Weimaraner | 0.38 | (-0.99, 1.75) | 0.59 | | | 1 |
| West Highland White Terrier | -0.84 | (-2.02, 0.35) | 0.17 | | | 1 |
| Whippet | -1.15 | (-3.48, 1.18) | 0.33 | | | 1 |
| Wirehaired Pointing Griffon | 3.80 | (-2.61, 10.21) | 0.24 | | | 1 |
| Yorkshire Terrier | -0.48 | (-1.92, 0.96) | 0.52 | | | 1 |

**Liver/Pancreas outcome: Logistic regression results**

Primary Liver/Pancreas Analysis: conditional logistic regression and the full analytic dataset, *n* = 24,238.

| **Explanatory variable** | **Estimated odds ratio** | **95% Confidence Interval** | ***p*** | **Degrees of freedom** |
| --- | --- | --- | --- | --- |
| feed once | 0.41 | (0.27, 0.61) | <0.001 | 1 |
| sex: female | 1.06 | (0.91, 1.23) | 0.47 | 1 |
| age |  |  | <0.001 | 4 |
| weight |  |  | <0.001 | 4 |

Secondary Liver/Pancreas Analysis: ordinary logistic regression using mixed-breed dogs and ten most populous breeds, *n* = 18,289.

| **Explanatory variable** | **Estimated odds ratio** | **95% Confidence Interval** | ***p*** | **Degrees of freedom** |
| --- | --- | --- | --- | --- |
| feed once | 0.25 | (0.12, 0.45) | <0.001 | 1 |
| sex: female | 1.16 | (0.96, 1.42) | 0.13 | 1 |
| age |  |  | <0.001 | 4 |
| weight |  |  | <0.001 | 4 |
| Australian Shepherd | 0.81 | (0.31, 1.71) | 0.61 | 1 |
| Beagle | 1.31 | (0.54, 2.69) | 0.51 | 1 |
| Border Collie | 1.25 | (0.52, 2.56) | 0.58 | 1 |
| Chihuahua | 1.53 | (0.70, 2.95) | 0.24 | 1 |
| Dachshund | 3.05 | (1.87, 4.80) | <0.001 | 1 |
| German Shepherd | 1.16 | (0.54, 2.21) | 0.67 | 1 |
| Golden Retriever | 0.64 | (0.35, 1.10) | 0.13 | 1 |
| Labrador Retriever | 0.95 | (0.61, 1.43) | 0.82 | 1 |
| Poodle (Standard)  (≥30 lbs) | 1.59 | (0.76, 2.95) | 0.18 | 1 |
| Pug | 0.79 | (0.24, 1.92) | 0.64 | 1 |

**Gastrointestinal outcome: Logistic regression results**

Primary Gastrointestinal Analysis: conditional logistic regression and the full analytic dataset, *n* = 24,238.

| **Explanatory variable** | **Estimated odds ratio** | **95% Confidence Interval** | ***p*** | **Degrees of freedom** |
| --- | --- | --- | --- | --- |
| feed once | 0.65 | (0.54, 0.77) | <0.001 | 1 |
| sex: female | 0.88 | (0.81, 0.95) | 0.0013 | 1 |
| age |  |  | <0.001 | 4 |
| weight |  |  | 0.068 | 4 |
| fatty acid | 1.13 | (1.02, 1.25) | 0.019 | 1 |

Secondary Gastrointestinal Analysis: ordinary logistic regression using mixed-breed dogs and ten most populous breeds, *n* = 18,289.

| **Explanatory variable** | **Estimated odds ratio** | **95% Confidence Interval** | ***p*** | **Degrees of freedom** |
| --- | --- | --- | --- | --- |
| feed once | 0.63 | (0.51, 0.78) | <0.001 | 1 |
| sex: female | 0.88 | (0.80, 0.98) | 0.017 | 1 |
| age |  |  | <0.001 | 4 |
| weight |  |  | 0.49 | 4 |
| fatty acid | 1.15 | (1.01, 1.30) | 0.035 | 1 |
| Australian Shepherd | 0.68 | (0.44, 1.03) | 0.071 | 1 |
| Beagle | 1.08 | (0.67, 1.74) | 0.76 | 1 |
| Border Collie | 1.12 | (0.74, 1.68) | 0.60 | 1 |
| Chihuahua | 1.09 | (0.69, 1.72) | 0.72 | 1 |
| Dachshund | 0.69 | (0.44, 1.03) | 0.10 | 1 |
| German Shepherd | 1.61 | (1.22, 2.12) | <0.001 | 1 |
| Golden Retriever | 0.84 | (0.67, 1.07) | 0.15 | 1 |
| Labrador Retriever | 0.85 | (0.68, 1.05) | 0.12 | 1 |
| Poodle (Standard)  (≥30 lbs) | 1.55 | (1.10, 2.17) | 0.012 | 1 |
| Pug | 1.96 | (1.32, 2.93) | <0.001 | 1 |

**Kidney/Urinary outcome: Logistic regression results**

Primary Kidney/Urinary Analysis: conditional logistic regression and the full analytic dataset, *n* = 24,238.

| **Explanatory variable** | **Estimated odds ratio** | **95% Confidence Interval** | ***p*** | **Degrees of freedom** |
| --- | --- | --- | --- | --- |
| feed once | 0.71 | (0.58, 0.88) | 0.0012 | 1 |
| sex: female | 3.09 | (2.77, 3.45) | <0.001 | 1 |
| age |  |  | <0.001 | 4 |
| weight |  |  | 0.11 | 4 |
| fatty acid | 1.22 | (1.09, 1.37) | <0.001 | 1 |

Secondary Kidney/Urinary Analysis: ordinary logistic regression using mixed-breed dogs and ten most populous breeds, *n* = 18,289.

| **Explanatory variable** | **Estimated odds ratio** | **95% Confidence Interval** | ***p*** | **Degrees of freedom** |
| --- | --- | --- | --- | --- |
| feed once | 0.67 | (0.51, 0.86) | 0.0024 | 1 |
| sex: female | 3.22 | (2.80, 3.71) | <0.001 | 1 |
| age |  |  | <0.001 | 4 |
| weight |  |  | 0.11 | 4 |
| fatty acid | 1.27 | (1.09, 1.46) | 0.0014 | 1 |
| Australian Shepherd | 0.90 | (0.55, 1.41) | 0.66 | 1 |
| Beagle | 1.35 | (0.81, 2.16) | 0.23 | 1 |
| Border Collie | 1.14 | (0.69, 1.79) | 0.59 | 1 |
| Chihuahua | 0.46 | (0.21, 0.87) | 0.028 | 1 |
| Dachshund | 0.84 | (0.52, 1.30) | 0.46 | 1 |
| German Shepherd | 1.12 | (0.75, 1.63) | 0.55 | 1 |
| Golden Retriever | 0.87 | (0.65, 1.15) | 0.33 | 1 |
| Labrador Retriever | 1.23 | (0.97, 1.54) | 0.079 | 1 |
| Poodle (Standard)  (≥30 lbs) | 0.61 | (0.32, 1.06) | 0.10 | 1 |
| Pug | 1.83 | (1.13, 2.86) | 0.010 | 1 |

**Orthopedic outcome: Logistic regression results**

Primary Orthopedic Analysis: conditional logistic regression and the full analytic dataset, *n* = 24,238.

| **Explanatory variable** | **Estimated odds ratio** | **95% Confidence Interval** | ***p*** | **Degrees of freedom** |
| --- | --- | --- | --- | --- |
| feed once | 0.78 | (0.69, 0.88) | <0.001 | 1 |
| sex: female | 1.01 | (0.96, 1.07) | 0.77 | 1 |
| age |  |  | <0.001 | 4 |
| weight |  |  | <0.001 | 4 |
| fatty acid | 1.60 | (1.49, 1.70) | <0.001 | 1 |

Secondary Orthopedic Analysis: ordinary logistic regression using mixed-breed dogs and ten most populous breeds, *n* = 18,289.

| **Explanatory variable** | **Estimated odds ratio** | **95% Confidence Interval** | ***p*** | **Degrees of freedom** |
| --- | --- | --- | --- | --- |
| feed once | 0.74 | (0.63, 0.87) | <0.001 | 1 |
| sex: female | 1.02 | (0.94, 1.11) | 0.63 | 1 |
| age |  |  | <0.001 | 4 |
| weight |  |  | <0.001 | 4 |
| fatty acid | 1.84 | (1.67, 2.03) | <0.001 | 1 |
| Australian Shepherd | 0.95 | (0.70, 1.29) | 0.75 | 1 |
| Beagle | 1.00 | (0.70, 1.45) | 0.98 | 1 |
| Border Collie | 1.10 | (0.79, 1.55) | 0.57 | 1 |
| Chihuahua | 1.04 | (0.72, 1.51) | 0.83 | 1 |
| Dachshund | 1.16 | (0.86, 1.56) | 0.34 | 1 |
| German Shepherd | 1.87 | (1.46, 2.41) | <0.001 | 1 |
| Golden Retriever | 1.08 | (0.89, 1.30) | 0.44 | 1 |
| Labrador Retriever | 1.61 | (1.38, 1.88) | <0.001 | 1 |
| Poodle (Standard)  (≥30 lbs) | 0.49 | (0.32, 0.74) | <0.001 | 1 |
| Pug | 1.10 | (0.76, 1.60) | 0.60 | 1 |

**Dental/Oral outcome: Logistic regression results**

Primary Dental/Oral Analysis: conditional logistic regression and the full analytic dataset, *n* = 24,238.

| **Explanatory variable** | **Estimated odds ratio** | **95% Confidence Interval** | ***p*** | **Degrees of freedom** |
| --- | --- | --- | --- | --- |
| feed once | 0.84 | (0.77, 0.92) | <0.001 | 1 |
| sex: female | 0.97 | (0.92, 1.01) | 0.17 | 1 |
| age |  |  | <0.001 | 4 |
| weight |  |  | <0.001 | 4 |

Secondary Dental/Oral Analysis: ordinary logistic regression using mixed-breed dogs and ten most populous breeds, *n* = 18,289.

| **Explanatory variable** | **Estimated odds ratio** | **95% Confidence Interval** | ***p*** | **Degrees of freedom** |
| --- | --- | --- | --- | --- |
| feed once | 0.77 | (0.67, 0.89) | <0.001 | 1 |
| sex: female | 0.95 | (0.88, 1.02) | 0.19 | 1 |
| age |  |  | <0.001 | 4 |
| weight |  |  | <0.001 | 4 |
| Australian Shepherd | 0.95 | (0.72, 1.26) | 0.73 | 1 |
| Beagle | 1.44 | (1.04, 1.99) | 0.027 | 1 |
| Border Collie | 1.23 | (0.90, 1.68) | 0.19 | 1 |
| Chihuahua | 3.52 | (2.60, 4.79) | <0.001 | 1 |
| Dachshund | 2.74 | (2.15, 3.49) | <0.001 | 1 |
| German Shepherd | 0.66 | (0.50, 0.87) | 0.0030 | 1 |
| Golden Retriever | 0.51 | (0.42, 0.63) | <0.001 | 1 |
| Labrador Retriever | 0.64 | (0.54, 0.75) | <0.001 | 1 |
| Poodle (Standard)  (≥30 lbs) | 0.92 | (0.68, 1.25) | 0.58 | 1 |
| Pug | 2.55 | (1.84, 3.52) | <0.001 | 1 |

**Cardiac outcome: Logistic regression results**

Primary Cardiac Analysis: conditional logistic regression and the full analytic dataset, n = 24,238.

| **Explanatory variable** | **Estimated odds ratio** | **95% Confidence Interval** | ***p*** | **Degrees of freedom** |
| --- | --- | --- | --- | --- |
| feed once | 0.86 | (0.70, 1.07) | 0.18 | 1 |
| sex: female | 0.85 | (0.76, 0.95) | 0.0054 | 1 |
| age |  |  | <0.001 | 4 |
| weight |  |  | <0.001 | 4 |
| fatty acid | 1.41 | (1.24, 1.61) | <0.001 | 1 |

Secondary Cardiac Analysis: ordinary logistic regression using mixed-breed dogs and ten most populous breeds, *n* = 18,289.

| **Explanatory variable** | **Estimated odds ratio** | **95% Confidence Interval** | ***p*** | **Degrees of freedom** |
| --- | --- | --- | --- | --- |
| feed once | 0.75 | (0.57, 1.00) | 0.048 | 1 |
| sex: female | 0.86 | (0.75, 1.00) | 0.049 | 1 |
| age |  |  | <0.001 | 4 |
| weight |  |  | <0.001 | 4 |
| fatty acid | 1.45 | (1.22, 1.72) | <0.001 | 1 |
| Australian Shepherd | 0.42 | (0.17, 1.04) | 0.062 | 1 |
| Beagle | 2.54 | (1.53, 4.20) | <0.001 | 1 |
| Border Collie | 1.00 | (0.50, 2.00) | 0.99 | 1 |
| Chihuahua | 5.76 | (3.89, 8.53) | <0.001 | 1 |
| Dachshund | 1.94 | (1.24, 3.02) | 0.0035 | 1 |
| German Shepherd | 0.68 | (0.33, 1.40) | 0.29 | 1 |
| Golden Retriever | 1.35 | (0.95, 1.93) | 0.097 | 1 |
| Labrador Retriever | 0.45 | (0.28, 0.72) | <0.001 | 1 |
| Poodle (Standard)  (≥30 lbs) | 1.18 | (0.63, 2.22) | 0.60 | 1 |
| Pug | 1.17 | (0.58, 2.37) | 0.66 | 1 |

**Cancer outcome: Logistic regression results**

Primary Cancer Analysis: conditional logistic regression and the full analytic dataset, *n* = 24,238.

| **Explanatory variable** | **Estimated odds ratio** | **95% Confidence Interval** | ***p*** | **Degrees of freedom** |
| --- | --- | --- | --- | --- |
| feed once | 0.90 | (0.75, 1.07) | 0.24 | 1 |
| sex: female | 1.01 | (0.92, 1.10) | 0.89 | 1 |
| age |  |  | <0.001 | 4 |
| weight |  |  | <0.001 | 4 |
| fatty acid | 1.29 | (1.16, 1.43) | <0.001 | 1 |

Secondary Cancer Analysis: ordinary logistic regression using mixed-breed dogs and ten most populous breeds, *n* = 18,289.

| **Explanatory variable** | **Estimated odds ratio** | **95% Confidence Interval** | ***p*** | **Degrees of freedom** |
| --- | --- | --- | --- | --- |
| feed once | 0.88 | (0.71, 1.10) | 0.26 | 1 |
| sex: female | 1.01 | (0.90, 1.13) | 0.91 | 1 |
| age |  |  | <0.001 | 4 |
| weight |  |  | <0.001 | 4 |
| fatty acid | 1.29 | (1.13, 1.47) | <0.001 | 1 |
| Australian Shepherd | 0.72 | (0.45, 1.14) | 0.16 | 1 |
| Beagle | 1.15 | (0.72, 1.84) | 0.55 | 1 |
| Border Collie | 0.50 | (0.27, 0.91) | 0.023 | 1 |
| Chihuahua | 0.34 | (0.17, 0.67) | 0.0018 | 1 |
| Dachshund | 0.41 | (0.24, 0.70) | <0.001 | 1 |
| German Shepherd | 0.51 | (0.30, 0.87) | 0.013 | 1 |
| Golden Retriever | 1.52 | (1.20, 1.91) | <0.001 | 1 |
| Labrador Retriever | 1.29 | (1.05, 1.60) | 0.018 | 1 |
| Poodle (Standard)  (≥30 lbs) | 1.01 | (0.65, 1.58) | 0.96 | 1 |
| Pug | 1.18 | (0.72, 1.92) | 0.51 | 1 |

**Neurological outcome: Logistic regression results**

Primary Neurological Analysis: conditional logistic regression and the full analytic dataset, *n* = 24,238.

| **Explanatory variable** | **Estimated odds ratio** | **95% Confidence Interval** | ***p*** | **Degrees of freedom** |
| --- | --- | --- | --- | --- |
| feed once | 0.90 | (0.71, 1.16) | 0.42 | 1 |
| sex: female | 0.79 | (0.70, 0.90) | <0.001 | 1 |
| age |  |  | <0.001 | 4 |
| weight |  |  | 0.14 | 4 |
| fatty acid | 1.28 | (1.11, 1.48) | <0.001 | 1 |

Secondary Neurological Analysis: ordinary logistic regression using mixed-breed dogs and ten most populous breeds, *n* = 18,289.

| **Explanatory variable** | **Estimated odds ratio** | **95% Confidence Interval** | ***p*** | **Degrees of freedom** |
| --- | --- | --- | --- | --- |
| feed once | 0.93 | (0.69, 1.25) | 0.63 | 1 |
| sex: female | 0.76 | (0.65, 0.89) | <0.001 | 1 |
| age |  |  | <0.001 | 4 |
| weight |  |  | 0.067 | 4 |
| fatty acid | 1.36 | (1.14, 1.63) | <0.001 | 1 |
| Australian Shepherd | 1.27 | (0.74, 2.18) | 0.39 | 1 |
| Beagle | 2.69 | (1.65, 4.41) | <0.001 | 1 |
| Border Collie | 1.07 | (0.54, 2.08) | 0.85 | 1 |
| Chihuahua | 0.91 | (0.45, 1.87) | 0.81 | 1 |
| Dachshund | 1.27 | (0.76, 2.13) | 0.37 | 1 |
| German Shepherd | 1.69 | (1.04, 2.75) | 0.035 | 1 |
| Golden Retriever | 1.49 | (1.06, 2.08) | 0.020 | 1 |
| Labrador Retriever | 1.23 | (0.90, 1.69) | 0.20 | 1 |
| Poodle (Standard)  (≥30 lbs) | 1.30 | (0.70, 2.40) | 0.41 | 1 |
| Pug | 2.31 | (1.39, 3.86) | 0.0013 | 1 |

**Skin outcome: Logistic regression results**

Primary Skin Analysis: conditional logistic regression and the full analytic dataset, *n* =24,238.

| **Explanatory variable** | **Estimated odds ratio** | **95% Confidence Interval** | ***p*** | **Degrees of freedom** |
| --- | --- | --- | --- | --- |
| feed once | 0.94 | (0.85, 1.04) | 0.22 | 1 |
| sex: female | 0.91 | (0.86, 0.95) | <0.001 | 1 |
| age |  |  | <0.001 | 4 |
| weight |  |  | <0.001 | 4 |
| fatty acid | 1.42 | (1.33, 1.51) | <0.001 | 1 |

Secondary Skin Analysis: ordinary logistic regression using mixed-breed dogs and ten most

populous breeds, *n* = 18,289.

| **Explanatory variable** | **Estimated odds ratio** | **95% Confidence Interval** | ***p*** | **Degrees of freedom** |
| --- | --- | --- | --- | --- |
| feed once | 0.95 | (0.83, 1.08) | 0.42 | 1 |
| sex: female | 0.88 | (0.82, 0.94) | <0.001 | 1 |
| age |  |  | <0.001 | 4 |
| weight |  |  | <0.001 | 4 |
| fatty acid | 1.53 | (1.40, 1.66) | <0.001 | 1 |
| Australian Shepherd | 0.70 | (0.55, 0.95) | 0.022 | 1 |
| Beagle | 0.71 | (0.49, 1.03) | 0.073 | 1 |
| Border Collie | 0.53 | (0.37, 0.75) | <0.001 | 1 |
| Chihuahua | 0.41 | (0.27, 0.62) | <0.001 | 1 |
| Dachshund | 0.66 | (0.49, 0.89) | 0.0071 | 1 |
| German Shepherd | 1.75 | (1.43, 2.14) | <0.001 | 1 |
| Golden Retriever | 1.24 | (1.07, 1.44) | 0.0041 | 1 |
| Labrador Retriever | 0.96 | (0.83, 1.10) | 0.54 | 1 |
| Poodle (Standard)  (≥30 lbs) | 0.78 | (0.58, 1.04) | 0.088 | 1 |
| Pug | 0.89 | (0.62, 1.28) | 0.53 | 1 |
