## Supplementary Information 4 for "Once-daily feeding is associated with better health in companion dogs: Results from the Dog Aging Project"

*Complete list of purebred dogs (n = 76) included in the CSLB analysis, with sample sizes.*

| **Breed** | **Sample size** |
| --- | --- |
| Labrador Retriever | 616 |
| Golden Retriever | 463 |
| German Shepherd | 192 |
| Dachshund | 159 |
| Australian Shepherd | 149 |
| Poodle (Standard) (≥30 lbs) | 130 |
| Border Collie | 124 |
| Chihuahua | 107 |
| Beagle | 105 |
| Shih Tzu | 93 |
| Yorkshire Terrier | 89 |
| Havanese | 82 |
| Miniature Schnauzer | 80 |
| Pug | 76 |
| Shetland Sheepdog | 76 |
| West Highland White Terrier | 75 |
| Cavalier King Charles Spaniel | 74 |
| Greyhound | 73 |
| Boston Terrier | 72 |
| Jack Russell Terrier | 69 |
| Cocker Spaniel | 64 |
| Pembroke Welsh Corgi | 62 |
| Boxer | 61 |
| English Springer Spaniel | 57 |
| German Shorthaired Pointer | 56 |
| Siberian Husky | 53 |
| Australian Cattle Dog | 51 |
| Poodle (Standard) (<30 lbs) | 50 |
| American Pitbull Terrier | 46 |
| Great Dane | 46 |
| Pomeranian | 45 |
| Cairn Terrier | 44 |
| Doberman Pinscher | 43 |
| Poodle (Toy) | 42 |
| Brittany | 39 |
| Bichon Frise | 38 |
| Soft Coated Wheaten Terrier | 37 |
| Weimaraner | 34 |
| Newfoundland | 33 |
| Maltese | 32 |
| Papillon | 31 |
| Rat Terrier | 31 |
| American Staffordshire Terrier | 30 |
| Bulldog | 29 |
| Collie | 29 |
| French Bulldog | 29 |
| Great Pyrenees | 28 |
| Vizsla | 28 |
| Portuguese Water Dog | 27 |
| Rottweiler | 27 |
| Bernese Mountain Dog | 26 |
| Basset Hound | 25 |
| border Terrier | 25 |
| Miniature Pinscher | 25 |
| Chesapeake Bay Retriever | 23 |
| English Setter | 23 |
| Rhodesian Ridgeback | 20 |
| Miniature American Shepherd | 19 |
| Scottish Terrier | 19 |
| Airedale Terrier | 18 |
| Dalmatian | 18 |
| Italian Greyhound | 18 |
| Shiba Inu | 18 |
| American Eskimo Dog | 17 |
| Samoyed | 17 |
| Alaskan Malamute | 16 |
| Whippet | 16 |
| Keeshond | 15 |
| Lhasa Apso | 15 |
| Bull Terrier | 14 |
| English Cocker Spaniel | 14 |
| Pekingese | 14 |
| Parson Russell Terrier | 13 |
| Carolina Dog | 12 |
| Catahoula Leopard Dog | 12 |
| Wirehaired Pointing Griffon | 11 |
