## Supplementary Information 5 for "Once-daily feeding is associated with better health in companion dogs: Results from the Dog Aging Project"

*Complete list of purebred dogs (n = 100) included in analysis of health conditions, with sample sizes.*

| **Breed** | **Sample size** |
| --- | --- |
| Labrador Retriever | 1467 |
| Golden Retriever | 1199 |
| German Shepherd | 502 |
| Australian Shepherd | 373 |
| Poodle (Standard) (≥30 lbs) | 306 |
| Dachshund | 305 |
| Border Collie | 257 |
| Chihuahua | 205 |
| Beagle | 187 |
| Pug | 180 |
| Miniature Schnauzer | 177 |
| Pembroke Welsh Corgi | 175 |
| Shih Tzu | 175 |
| Boxer | 173 |
| Havanese | 168 |
| Yorkshire Terrier | 160 |
| Cavalier King Charles Spaniel | 159 |
| Greyhound | 155 |
| Siberian Husky | 141 |
| Boston Terrier | 140 |
| Shetland Sheepdog | 137 |
| English Springer Spaniel | 135 |
| Australian Cattle Dog | 133 |
| Great Dane | 132 |
| Cocker Spaniel | 125 |
| Doberman Pinscher | 122 |
| German Shorthaired Pointer | 122 |
| French Bulldog | 121 |
| Jack Russell Terrier | 119 |
| West Highland White Terrier | 119 |
| American Pitbull Terrier | 101 |
| Poodle (Standard) (<30 lbs) | 98 |
| Pomeranian | 88 |
| Bernese Mountain Dog | 85 |
| Newfoundland | 84 |
| Collie | 83 |
| Soft Coated Wheaten Terrier | 83 |
| Great Pyrenees | 80 |
| Poodle (Toy) | 80 |
| Bichon Frise | 77 |
| Basset Hound | 76 |
| Brittany | 74 |
| Maltese | 74 |
| Vizsla | 74 |
| Rottweiler | 70 |
| Bulldog | 68 |
| American Staffordshire Terrier | 67 |
| Cairn Terrier | 67 |
| Weimaraner | 67 |
| Miniature American Shepherd | 65 |
| Shiba Inu | 61 |
| Rat Terrier | 57 |
| Rhodesian Ridgeback | 52 |
| Portuguese Water Dog | 51 |
| English Setter | 50 |
| Scottish Terrier | 45 |
| Cardigan Welsh Corgi | 44 |
| Miniature Pinscher | 43 |
| Papillon | 43 |
| Chesapeake Bay Retriever | 42 |
| Dalmatian | 41 |
| Airedale Terrier | 40 |
| Alaskan Malamute | 40 |
| St. Bernard | 40 |
| Belgian Malinois | 38 |
| Italian Greyhound | 38 |
| border Terrier | 37 |
| Whippet | 35 |
| American Eskimo Dog | 34 |
| Coton De Tulear | 34 |
| Keeshond | 34 |
| English Cocker Spaniel | 32 |
| Catahoula Leopard Dog | 31 |
| Samoyed | 31 |
| Mastiff | 30 |
| Akita | 25 |
| Lhasa Apso | 25 |
| Pekingese | 25 |
| Wirehaired Pointing Griffon | 25 |
| Welsh Terrier | 24 |
| Basenji | 22 |
| Carolina Dog | 22 |
| Nova Scotia Duck Tolling Retriever | 22 |
| Bouvier des Flandres | 21 |
| Bull Terrier | 21 |
| Parson Russell Terrier | 21 |
| Norwich Terrier | 20 |
| Irish Wolfhound | 19 |
| Boykin Spaniel | 18 |
| Schipperke | 18 |
| Chinese Shar-Pei | 17 |
| German Wirehaired Pointer | 17 |
| Tibetan Terrier | 17 |
| Chinese Crested | 16 |
| Wire Fox Terrier | 16 |
| English Shepherd | 14 |
| Pointer | 14 |
| Standard Schnauzer | 14 |
| Bloodhound | 13 |
| Gordon Setter | 11 |
